# Recovering short DNA fragments from minerals and marine sediments: a comparative study evaluating lysis and isolation approaches

**DOI:** 10.1101/2023.12.09.570911

**Authors:** Darjan Gande, Christiane Hassenrück, Marina Žure, Tim Richter-Heitmann, Eske Willerslev, Michael W. Friedrich

**Author notes:** **Corresponding authors:** Darjan Gande; and Michael W. Friedrich.

## Abstract

Marine sediments as excellent climate archives, contain among other biomolecules substantial amounts of extracellular DNA. Through mechanisms of binding to various minerals, some of the DNA stays protected from degradation and remains preserved. While this pool of DNA represents genomic ecosystem fingerprints spanning over millions of years, the capability of current DNA extraction methods in recovering mineral-bound DNA remains poorly understood. We evaluated current sedimentary DNA extraction approaches and their ability to desorb and extract short DNA fragments from pure clay and quartz minerals as well as from different types of marine sediments. We separately investigated lysis (DNA release) and isolation steps (purification of DNA) comparing five different types of lysis buffers across two commonly used DNA isolation approaches: silica magnetic beads and liquid-phase organic extraction and purification. The choice of lysis buffer significantly impacted the amount of recovered mineral-bound DNA and facilitated selective desorption of DNA fragments. High molarity EDTA and phosphate lysis buffers recovered on average an order of magnitude more DNA from clay than other tested buffers, while both isolation approaches recovered comparable amounts of DNA. However, organic extraction caused an inhibitory effect in subsequent downstream applications (e.g., PCR), across all assessed sediment DNA extracts while silica magnetic beads induced inhibition only in half of the tested DNA extracts. Furthermore, we demonstrate that both, the isolation approach and the lysis buffer play a decisive role in successful library preparation and that lysis buffer choice impacted the final library fragment distribution. With this study, we underscore the critical importance of lysis buffer selection to maximize the recovery of mineral-bound DNA in sedimentary DNA extractions and show its profound impact on recovered fragment lengths, a crucial factor alongside existing isolation approaches in facilitating high-quality DNA extracts for downstream analysis related to ancient environmental (aeDNA) research.

## 1. Introduction

In recent years the field of ancient environmental (aeDNA) reached remarkable progress in unravelling unprecedented details of the structure of ancient ecosystems and their diversity dynamics in regard to climate perturbations on millennial and even geological time scales (Wang et al. 2021, Alsos et al. 2022, Armbrecht et al. 2022, Kjær et al. 2022). Driven by continuously developing sequencing technologies and genetic techniques, these aeDNA analyses from aquatic and terrestrial deposits not only capture past ecosystem biodiversity but also hold increasingly important potential for understanding future ecosystem structure (Crump 2021).

To date, studies employing aeDNA as a climate and biodiversity proxy in sediments mostly focused on lacustrine deposits (Nguyen et al. 2023), with a marked increase in aeDNA research on marine deposits observed recently (e.g., De Schepper et al. 2019, Zimmermann et al. 2020, Zimmermann et al. 2021, Selway et al. 2022), including latest reports of 1-million old diatom DNA isolated from marine sediment core (Armbrecht et al. 2022) and a record 2-million old DNA from Kap København formation, which in part includes old marine sediments (Kjær et al. 2022). Although fossilizing taxa (e.g., foraminifera, diatoms, coccolithophores) offer a well-established proxy for marine paleoclimate reconstructions, aeDNA adds to a deeper understanding of past ecosystem structure capturing diversity beyond the fossilized repository and often complementing it (Lejzerowicz et al. 2013, Parducci et al. 2019). Combined with existing climate proxies, marine paleoenvironmental reconstructions using aeDNA offer a window of opportunity for understanding the evolution and dynamics of marine ecosystems over geological timescales on a local and global scale, especially in view of ongoing climate change (Coolen et al. 2006, De Schepper et al. 2019, Epp 2019, Armbrecht et al. 2022).

In general, ancient DNA is typically highly fragmented to sizes rarely exceeding < 100 bp (Gamba et al. 2016) and exhibits unique damage characteristics of cytosine to uracil deamination at the overhanging ends of the molecule (Briggs et al. 2007, Orlando et al. 2021). Preservation of these degraded DNA fragments from various organisms through millions of years is likely facilitated by stable cold temperatures and binding of DNA to mineral surfaces, where it remains shielded from complete enzymatic degradation (Romanowski et al. 1991, Willerslev et al. 2004, Kjær et al. 2022).

In fact, binding of DNA to mineral components of soils and sediments is well characterized and has been studied in the past (e.g., Lorenz and Wackernagel 1987, Romanowski et al. 1991, Khanna and Stotzky 1992, Lorenz and Wackernagel 1992, Paget et al. 1992, Cai et al. 2006). DNA binds rapidly and forms stable complexes with minerals such as sand and clay, with the latter exhibiting approximately 700-fold higher binding capacity than sand (Lorenz and Wackernagel 1994). In addition, clays such as montmorillonite (smectite group) and kaolinite also exhibit higher binding capacity for low molecular weight DNA as opposed to high molecular weight DNA (Pietramellara et al. 2001).

These interactions are mainly facilitated by electrostatic forces, ligand exchange, hydrogen bonding and cation bridging with clay mineral structure, surface topography, and surface charge directly affecting the amount of adsorbed DNA as well as its adsorption behavior and preservation (Yu et al. 2013, Freeman et al. 2023). The adsorption of DNA to these clay minerals is thus governed by ionic conditions (i.e., concentration, valency, type) and pH (Yu et al. 2013 and references therein). Importantly, such mineral-bound DNA exhibits resistance towards direct enzymatic degradation (Torti et al. 2015); for example, 10-fold and 100-1000-fold more DNase enzyme was required to achieve similar degradation of DNA bound to clay and sand, respectively, in comparison to dissolved DNA (Romanowski et al. 1991, Khanna and Stotzky 1992).

With over 90% of DNA in deep-sea sediments present outside of cells (DelĺAnno and Danovaro 2005) and an estimated 50-95% of that being bound to the sediment matrix (Torti et al. 2015 and references therein), this pool of protected DNA represents a large reservoir of available genetic information about past biodiversity. In turn, this underlines the significance of marine sediments as global climate archives for studying past ecosystem structure and climate impacts on it, for which choosing the most suitable extraction approach to maximize the recovery of mineral-bound DNA is paramount.

In the past decade, studies focusing on extraction of aeDNA from different sediments and soils used a broad range of methods to retrieve aeDNA from sediment material differing in the lysis step (release of DNA into the solution) and the subsequent DNA isolation steps (purification of DNA). Commercial soil extraction kits have been utilized most often (e.g., Coolen et al. 2013, Lejzerowicz et al. 2013, Capo et al. 2015, Capo et al. 2016), with some studies modifying the kit protocol’s lysis conditions (Stoof-Leichsenring et al. 2015, Zimmermann et al. 2017, Epp et al. 2019, Voldstad et al. 2020, Zimmermann et al. 2020). More recently, lysis conditions based on ∼0.5M EDTA otherwise regularly used in bone and tooth DNA extractions (Dabney et al. 2013, Korlević et al. 2015, Margaryan et al. 2020) also proved successful in sedimentary aeDNA extractions, combined either with silica-spin columns (Slon et al. 2017), liquid silica (Armbrecht et al. 2020) or silica magnetic beads (Rohland et al. 2018) as final DNA isolation steps. Less often used were phosphate-based lysis buffers - combined either with silica-spin columns (Lammers et al. 2021) or liquid-phase organic extraction with phenol-chloroform-isoamylalcohol (P/C/I) and ultrafiltration-based up-concentration (Orsi et al. 2017, More et al. 2018) to isolate the DNA. Multicomponent “Bulat” lysis buffer (Bulat Sergey et al. 2000) containing Tris-HCl, EDTA and NaCl, combined with a similar isolation approach utilizing P/C/I and ultrafiltration was used in several studies albeit with slight modifications (Pedersen et al. 2016, Parducci et al. 2019) and has recently also facilitated the isolation of 2-million-old DNA (Kjær et al. 2022).

Several above-mentioned DNA isolation approaches have been extensively evaluated in their impact on fragment length recovery considering the fragmented nature of ancient DNA (e.g., Dabney et al. 2013, Allentoft et al. 2015, Rohland et al. 2018). Most notably, silica spin columns were shown to favor recovery of longer DNA fragments over short ones, the latter more likely representing aeDNA (Rohland et al. 2018). This bias, however, has been shown to be surmountable by the use of silica magnetic beads (Rohland et al. 2018) and – as more recently reported – ethanol-based precipitations using linear polyacrylamide (Maixner et al. 2021) facilitating reliable recovery of short DNA fragments from the sediment matrix.

While the choice of the DNA isolation approach is an important factor to consider in the retrieval of short DNA fragments, performance and impact of the initial part of DNA extraction, the lysis, in desorbing mineral-bound DNA fragments into solution remains poorly understood. To date, high molarity phosphate and EDTA solutions were shown to outperform soil extraction kit buffers and the “Bulat” buffer in recovering short DNA fragments especially from clay minerals (Direito et al. 2012, Slon et al. 2017). This is expected as EDTA is a chelator of divalent cations and by sequestering them can disrupt the DNA-cation-mineral bridge. However, the complexity of DNA pools in marine sediments suggested by Torti et al. (2015) implies that lysis conditions should target not only mineral-bound DNA but also DNA residing in dormant/deceased forms of planktonic organisms (Armbrecht et al. 2019). Such an approach has recently been shown as the most comprehensive strategy in detecting eukaryotes in marine sediments (Armbrecht et al. 2020) and thus, underlines the importance of using sophisticated lysis conditions to retrieve aeDNA residing in multiple DNA pools.

In light of the latest advances in the field of aeDNA research, demonstrating that DNA preservation on million-year scales is strongly linked to mineral binding (Kjær et al. 2022), possibilities of studying such old deposits began to widen. Although to date, the exact proportion of mineral-bound DNA in sedimentary deposits is not well constrained, currently used sedimentary DNA extraction approaches such as the ones described above, have until now not been systematically investigated in their capabilities and shortcomings in regard to desorbing mineral-bound DNA.

In this study we aimed to address this gap and provide more compelling details on the effect of the initial lysis step of multiple DNA extraction protocols on desorption and retrieval of mineral-bound DNA. We systematically evaluated five different lysis conditions previously used in various aeDNA and other sedimentary DNA extractions (Direito et al. 2012, Lever et al. 2015, Pedersen et al. 2016, Rohland et al. 2018, Epp et al. 2019) and assessed their ability to desorb and extract short DNA fragments from two pure primary clay minerals present in marine sediments (bentonite (smectite group) and kaolinite), pure sea sand (quartz) and different types of natural marine sediments varying in organic matter content, general lithology and depth/age. In addition, we compared lysis conditions across two DNA isolation approaches mentioned above: silica magnetic beads and liquid-phase organic extraction using phenol/chloroform/isoamyl alcohol and precipitation with ethanol and linear polyacrylamide.

## 2. Methods

### 2.1. Binding short DNA fragments to the minerals

To facilitate DNA binding to the mineral structure in conditions similar to the ocean floor-seawater interface, we mixed 4 µL (2 µg) of GeneRuler Ultra Low DNA Ladder (10-300 bp; Thermo Fischer Scientific, USA) with 996 µL sterile-filtered artificial sea water (composition [L^−1^]: 26.4 g NaCl, 11.2 g MgCl_2_⋅6H_2_O, 1.5 g CaCl_2_⋅2H_2_O and 0.7 g KCl) and 0.15 g of the given mineral in a sterile 2.0 mL screw-cap tube – a modified approach originally described by Slon et al. (2017). The tubes were incubated for 24 hours at room temperature under gentle horizontal mixing to allow DNA to bind to the matrix and then centrifuged at 20817 × *g* for 10 minutes to separate the mineral and liquid fraction. The supernatant was removed and frozen (-20°C) for later isolation of un-bound DNA and the mineral pellet was subjected to DNA extraction.

### 2.2. Lysis – desorption of DNA from minerals

To each pellet, we first added ∼0.7 g of sterile 0.1 mm Zirconia beads (Carl Roth, Germany), followed by the addition of the lysis buffer: Rohland (Rohland et al. 2018), Direito (Direito et al. 2012), PowerSoil (Epp et al. 2019), Pedersen (Pedersen et al. 2016) and Lever (Lever et al. 2015) (see Table 1 for more details). Each sample was then homogenized on a FastPrep-24™ homogenizer (MP Biomedicals, USA) at 4.5 m/s for 20 seconds and incubated according to the protocol (Table 1) with modifications (see Supplementary Information, 1.). After centrifugation at 20817 × *g* for 10 minutes to remove all matrix particles the resulting lysate was transferred to a fresh 2.0 mL Lo-Bind tube (Eppendorf, Germany). DNA was isolated from the lysate using two approaches: silica magnetic beads and phenol-chloroform-isoamyl alcohol (PCI/CI) followed by ethanol precipitation and linear polyacrylamide (LPA).

**Table 1.** Buffer composition and lysis conditions used in this study.

| Reference | Buffer name | Buffer Composition | Volume per sample + additions | Incubation |
| --- | --- | --- | --- | --- |
| Rohland, et al. 2018 | Rohland | 450 mM EDTA<br>0.05% Tween20 | 1 mL buffer + 250 µg Proteinase K | 16h at 37°C, mixing |
| Direito, et al. 2012 | Direito | 1M Phosphate buffer / 15% Ethanol | 1 mL buffer + 60 µL Qiagen MagAttract PowerSoil kit CD1 solution | 40 min at 80°C, mixing |
| Epp, et al. 2019 | PowerSoil | <u>Qiagen MagAttract PowerSoil kit (2020)</u><br>CD1 + CD2 solution | 800 µL CD1 + 100 µg Proteinase K<br>+<br>200 µL CD2 solution (*subsequent step*) | 2h at 45°C, mixing |
| Pedersen, et al. 2016 | Pedersen | Stock buffer:<br>50 mM Tris-HCl,<br>20 mM EDTA,<br>150 mM NaCl,<br>2% N-laurylsarcosine<br><u>Final buffer:</u><br>• 1 mL 2-Mercaptoethanol,<br>• 1.5 mL 1M dithiothreitol<br>• 27.5 mL stock buffer | 1 mL buffer + 115 µg Proteinase K | 16h at 37°C, mixing |
| Lever, et al. 2015 | Lever | 30 mM Tris-HCl<br>30 mM EDTA<br>1% Triton-X-100<br><u>800 mM Guanidinium Hydrochloride</u><br>Adjust pH to 10 | 200 µL 10 mM Na-hexametaphosphate<br>+<br>1 mL buffer | 1h at 50 °C, mixing |

### 2.3. DNA isolation using silica magnetic beads

For DNA isolation using silica magnetic beads we followed the protocol described Rohland et al. (2018) with modifications (Supplementary Information, 2.). Each extract was first concentrated using Amicon® Ultra 0.5 mL, 10 kDa centrifugal filters (MERCK, Germany) with 2x 10-minute centrifugations at 15.294 × *g*, washed with 450 µL of EB buffer (10 mM Tris-Cl, pH 8.5; Qiagen, Germany) and centrifuged again at 15.294 × *g* for 10 minutes to a final volume of 120 µL. The retentate was transferred into a clean 2.0 mL Lo-Bind tube (Eppendorf, Germany) and mixed with 1680 µL of modified PB buffer (Qiagen, Germany) at a ratio of 15:1 (see Supplementary Information, 2. for more details) and 10 µL silica magnetic beads (G-Biosciences, USA). The mixture was briefly vortexed and incubated for 20 minutes at room temperature while shaking at 1000 rpm to facilitate DNA binding. Beads were collected on a magnet, the supernatant was removed and beads were washed two times using 500 µL PE wash buffer (Qiagen, Germany) before air-drying. DNA bound to the beads was eluted using 30 µL EB buffer (Qiagen, Germany), incubated for 10 minutes under gentle mixing at room temperature after which the beads were collected on a magnet and the eluate was transferred into a clean 1.5 mL Lo-Bind tube (Eppendorf, Germany).

### 2.4. DNA isolation using phenol-chloroform-isoamyl alcohol and ethanol precipitation

Liquid-phase organic extraction of DNA using PCI/CI was based on previous studies (Lever et al. 2015, Maixner et al. 2021) albeit with modifications. Each lysate was mixed with one volume of cooled (4°C) phenol-chloroform-isoamyl alcohol (P/C/I) (25:24:1, vol/vol/vol) (Carl Roth, Germany) followed by centrifugation at 20817 × *g* and 4°C for 5 minutes after which the upper aqueous part was transferred to a new tube. The process was repeated with one volume of cooled (4°C) chloroform-isoamyl alcohol (C/I) (24:1, vol/vol) (Carl Roth, Germany). In case of the Direito lysis buffer, the centrifugation was performed at 35°C to avoid phosphate precipitation from the buffer. Processed extracts, except those obtained using lysis conditions described in Lever et al. (2015) and Pedersen et al. (2016), were additionally de-salted using Amicon® Ultra 0.5 mL, 10 kDa centrifugal filters (MERCK, Germany) as described above, followed by a final volume adjustment to 500 µL with EB buffer (Qiagen, Germany). This step was necessary to enable effective DNA precipitation while avoiding salt co-precipitation due to lysis buffers containing high EDTA and phosphate concentrations.

Finally, DNA was precipitated with the addition of 4 µL of 25 mg/mL GeneElute Linear Polyacrylamide (SigmaAldrich, USA) and 2.5-3 volumes of 100% ethanol to each extract. After 30-minute incubation at room temperature, tubes were centrifuged at 4°C for 45 minutes and 20817 × *g*. Resulting DNA pellets were washed with 1.0 mL 70% ice-cold ethanol and re-suspended in 30 µL of EB buffer (Qiagen, Germany) after drying.

DNA concentrations of each extract were determined using Quant-iT™ PicoGreen™ dsDNA Assay (Invitrogen, USA) and the total DNA amount was calculated per total volume of the final extract. The extracts were visualized on a 5% agarose gel run at 5 V/cm for one hour in 1X TBE buffer and stained with ethidium bromide.

In summary, we tested five different lysis conditions (Rohland, Direito, PowerSoil, Pedersen and Lever) over two different isolation approaches (silica magnetic beads and organic extraction using PCI/CI with precipitation) for three different minerals (bentonite, kaolinite and sea-sand). Each combination of lysis buffer, mineral, and isolation approach had a minimum of four replicates.

### 2.5. Extraction of DNA from sediment samples

We further assessed the performance of the lysis buffers by Rohland et al. (2018) and Direito et al. (2012) on five biogenically different sediment samples (hereafter referred to with the given abbreviation): North Pacific ocean siliciclastic silicious (N-PAC) and siliciclastic carbonaceous sediment (MID-PAC), Black Sea organic sapropel (BS-SAP) and terrigenous/lacustrine mud (BS-LAC) and coastal diatom rich South Georgia Island sediment (SG-CLY) (see Table S4 for further details).

Performance of the two buffers and isolation approaches was evaluated based on their ability to extract short DNA fragments from the sediment matrix and overall extract quality in terms of inhibition of downstream processes.

Prior to extractions, frozen sediment was thawed, homogenized and partitioned into 0.3 g aliquots in 2.0 mL screw-cap tubes under a laminar flow hood with precautions taken to prevent outside contamination and cross-contamination by using sterile consumables and wiping working surfaces with a 1-2% hypochlorite solution. Aliquots were kept frozen until extractions commenced. Sediment samples were processed as described under 2.2. using the two above-mentioned lysis buffers and DNA isolated with the two isolation approaches described under 2.3. and 2.4. All extraction steps except PCI/CI addition and centrifugations were performed under a laminar flow hood. One extraction blank control (only lysis buffer) was included for each combination of lysis buffer and isolation approach.

To investigate the possible range of DNA fragment lengths extractable from the sediment matrix we spiked additional sediment aliquots (0.3 g) with DNA ladder as described under 2.1. and extracted the DNA as mentioned above for the un-spiked sediment samples.

All sediment DNA extractions (with and without a spike) were performed in duplicates.

#### 2.5.1. Evaluating PCR inhibition of sedimentary DNA extracts

We used bacterial 16S rRNA gene PCR spiked with *Escherichia coli* genomic DNA for evaluating the effect of co-extracted inhibitory substances such as phenol, ethanol, salts and humics in sedimentary DNA extracts on DNA-modifying enzyme reactions involved in downstream processing such as sequencing library preparation.

For this, two replicates of un-spiked sedimentary DNA extracts were pooled together first. A 25 µL PCR reaction was then set up for each sample, containing 0.5 µL dNTP mix (10 mM), 1.5 µL MgCl_2_ (25 mM), 2.5 µL 10X polymerase buffer, 0.25 µL bovine serum albumin (20 mg/mL), 0.75 µL each of 8-Forward and 338-Reverse primer (10 µM each), 0.25 µL Taq Polymerase (Applied Biosystems™), 2 µL DNA extract and 1 µL of *E. coli* genomic DNA (200 ng/µL). The reaction was run for 40 cycles with the following settings: initial denaturation (95°C, 5 min), denaturation (95°C, 30 sec), annealing (58°C, 30 sec), elongation (72°C, 30 sec) and final elongation (72°C, 5 min). Resulting fragments expected at a length of 330 bp were visualized on a 2% agarose gel run at 5 V/cm for 1 hour in 1X TAE buffer stained with ethidium bromide.

Sediment extracts for which the PCR reaction did not yield a visible band compared to the positive control (*E. coli* genomic DNA without sedimentary DNA extract) were considered inhibited and were subsequently purified using the One-Step Inhibitor Removal kit (Zymo Research Europe, Germany) following the manufacturer’s instructions. Inhibition PCR reactions were repeated for the purified extracts and products again visualized by analytical agarose gel electrophoresis as described above.

### 2.6. Preparation of sequencing libraries and fragment analysis

As previously suggested, extracts containing low to undetectable concentrations of DNA are best characterized when converted into sequencing libraries (Rohland et al. 2018). We therefore used un-spiked sediment DNA extracts to build double-stranded libraries for subsequent analysis of fragment distribution and final comparison of the performance of the two lysis and isolation approaches. Due to the differences in DNA concentrations in pooled extracts across the sediment samples, we adjusted the input volume for each extract in the first step of library preparation (end-repair) and consequently used proportionally more enzyme and buffer. We limited the total input of DNA for library preparation to 100 ng or less. Details on final concentrations, amount of input DNA for library preparation and inhibition are shown in Table SE1. Sequencing libraries were prepared and later purified following the protocol published by (Meyer and Kircher 2010) and used previously (Pedersen et al. 2016, Wang et al. 2021, Kjær et al. 2022) with 10.5 µL prepared library used for indexing PCR (iPCR). Number of cycles for indexing PCR were adjusted for each sample according to a screening qPCR to avoid possible library overamplification (Table SE1).

Fragment distribution in resulting libraries was assessed for each library separately by means of capillary electrophoresis using a Fragment Analyzer (Agilent Technologies, USA). Based on the Fragment Analyzer results we later calculated the molarity ratio between fragment ranges with sufficiently long insert (157 – 500 bp) and fragment range corresponding to adapter dimers and/or short inserts (25 – 157 bp) to evaluate overall library quality. With adapter dimers expected at a length of ∼127 bp, the lower threshold for sufficiently long fragment of 157 bp would represent a ∼30 bp long insert. Thus, the higher the ratio, the higher the molarity of sufficiently long inserts in the final library.

Furthermore, to understand the impact of lysis buffer and isolation approach on fragment length distribution in prepared libraries we evaluated molarities of selected fragment ranges. Using the ProSize Data Analysis Software (Agilent Technologies, USA) we divided the chromatograms into ranges of 157-177 bp, 177-202 bp, 202-227 bp, 227-277 bp, 277-327 bp and 327-427 bp, and retrieved the molarity (nM) for each range. When subtracting the adapter sequence at ∼127 bp, these ranges would correspond to approximately 30-50 bp, 50-75 bp, 75-100 bp, 100-150 bp, 150-200 bp and 200-300 bp long inserts.

### 2.7. Statistical analysis of mineral-bound DNA extractions and sediment library fragment size distributions

All statistical analyses were conducted using R (v. 4.2.3) and RStudio (v. 2023.03.1+446) (Posit Team 2023, R Core Team 2023). Our experimental design was initially balanced but during the extractions we noticed that for some lysis buffer / mineral combinations, total extracted DNA values of replicates exhibited strong variation (Table S2). We thus included more replicates and used a non-parametric Scheirer–Ray–Hare test (Scheirer et al. 1976) within the R package rcompanion (v. 2.4.30) (Mangiafico 2023) to test for the effect of lysis buffer, isolation approach and their interaction on total extracted DNA for each mineral separately (a two-way analysis using type-II sum-of-squares). A post-hoc analysis was subsequently carried out using Dunn tests (Dunn 1961) within R package FSA (v. 0.9.4) (Dinno 2022) for the lysis buffer effect for each mineral type. All p-values were adjusted using the “holm” method and differences in medians between different buffers summarized using compact letter display based on a family-wise statistical significance level at 0.05 (Piepho 2004).

With the sequencing libraries prepared from natural sediments we explored the effect of lysis buffer on relative and absolute molarities across selected fragment ranges. We used linear mixed-effects models within the R package lme4 (v. 1.1-33) (Bates et al. 2015) with lysis buffer as fixed factor and sediment type as random effects variable. Subsequent post-hoc pairwise comparisons between lysis buffers were performed with Tukey’s contrasts using R package emmeans (v. 1.8.8) at a statistical significance of 0.05 (Searle et al. 1980).

## 3. Results and discussion

### 3.1. Total DNA recovered from minerals

In this experiment, we aimed to evaluate the impact of lysis buffer on the total amount of desorbed DNA from mineral structures over two DNA isolation approaches. We detected significant differences between lysis buffers on total extracted DNA for all three mineral types (Scheirer-Ray-Hare test, *P* < 0.05), while no significant effect was found for either the isolation approach or the interaction of lysis buffer and isolation approach (Scheirer-Ray-Hare test, *P* > 0.05) (see Table S3 for more details).

Subsequent post-hoc tests of buffer effect for bentonite and kaolinite clays revealed that Rohland and Direito lysis buffers performed equally well, recovering on average an order of magnitude more DNA compared to the other three lysis buffers. While Pedersen and Lever lysis buffers consistently recovered the least DNA from bentonite, showing significant differences to Rohland and Direito buffers, the performance of the PowerSoil buffer was more heterogeneous and only significantly different from the Lever buffer (Figure 1A). In the case of kaolinite, however, PowerSoil and Lever buffers recovered least DNA showing significant differences to the best-performing Rohland and Direito buffers, while the Pedersen buffer achieved intermediate DNA recovery that was only significantly lower than that of the Direito buffer (Figure 1C).

**Figure 1 A-C.**
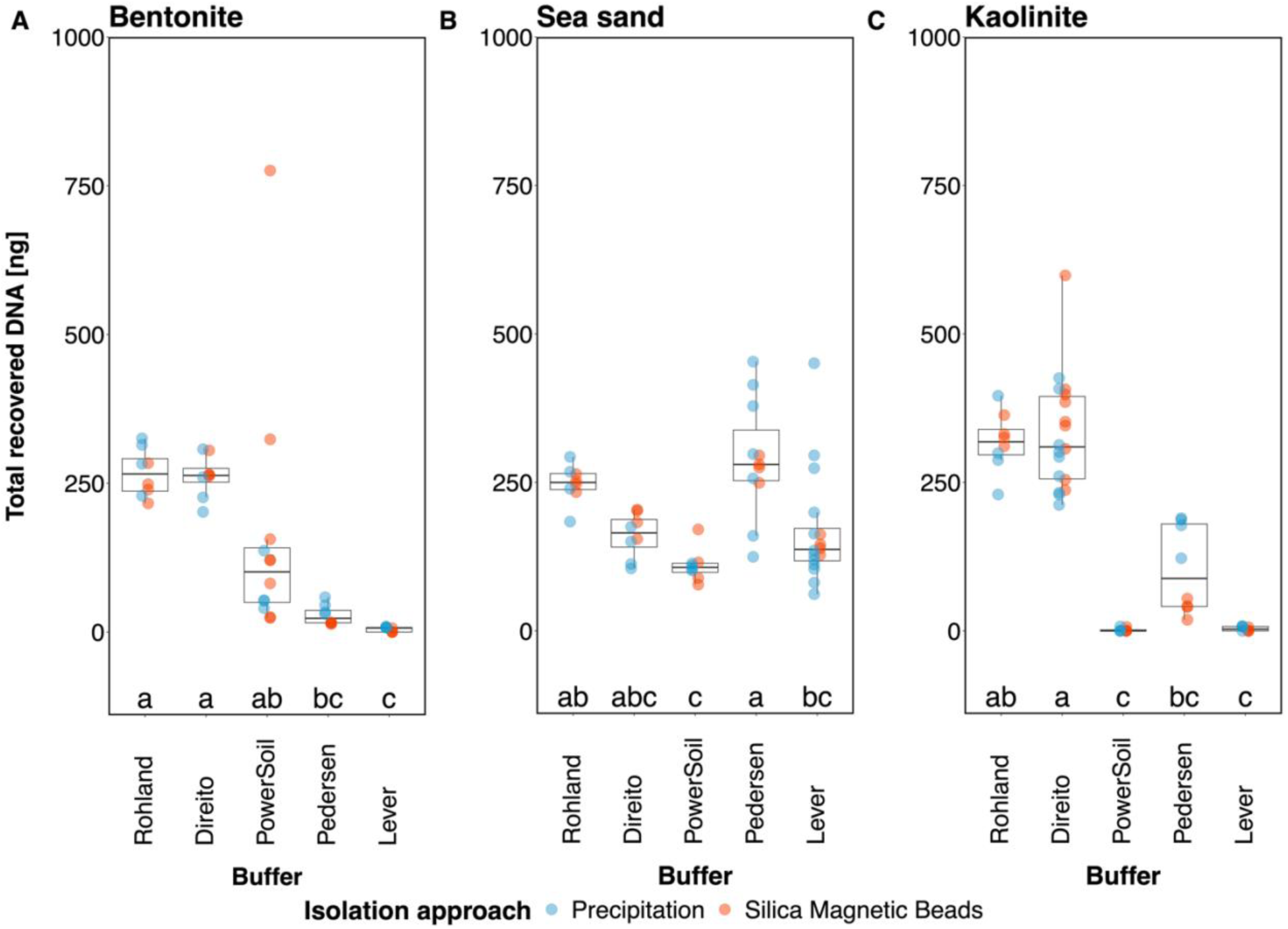
Total recovered DNA (ng) from A) bentonite, B) sea-sand and C) kaolinite using five different lysis buffers and isolated with either silica magnetic beads or PCI/CI and precipitation. Letters depict results of post-hoc tests conducted to compare the median values for the amount of recovered DNA. Grouping based on a statistical significance threshold at 0.05.

In contrast to the clay minerals, recovery of DNA from sea-sand appeared more similar across the different lysis buffers. Here, Pedersen buffer recovered the highest amounts of DNA on average, showing significant differences only to PowerSoil and Lever buffers. In addition, Direito and Rohland buffers exhibited similar performance to the Pedersen buffer with the latter also performing significantly better than the PowerSoil buffer (Figure 1B).

These inter- and intra-mineral buffer performance variations especially on clays can be explained by the interaction between the strength of the DNA-mineral bond and the composition of the lysis buffer. As summarized by Yu et al. (2013), depending on their type, clay minerals contain one or more adsorption sites - interlamellar channels and external surfaces such as broken edges or planar surface with which the DNA primarily interacts with, either through cation bridging or ligand exchange, hydrogen bonding and electrostatic forces respectively. Taking this into consideration it is reasonable to assume that the lysis buffer (depending on its composition) may therefore only achieve partial desorption of DNA from particular sites where it can successfully break the mineral-DNA bond. This can be observed clearly for the PowerSoil buffer (Qiagen, proprietary composition) which was able to desorb low amounts of DNA from bentonite, where DNA mainly adsorbs to planar surface though weaker interactions but no DNA from kaolinite, where DNA has been shown to mainly bind to the broken edges with strong interactions (see Yu et al. 2013 and references therein).

Furthermore, the total amount of recovered DNA from the minerals by different lysis conditions was rather low - less than 50% of the input even though the amount of unbound DNA after DNA adsorption to the mineral was negligible at an average 0.45 % (bentonite), 1.14 % (sea-sand) and 0.32 % (kaolinite) of total input DNA (see Supplementary Information 3.). We therefore investigated the capacity and constrains of the two DNA isolation approaches with DNA ladder only (no mineral involved) as described under 2.3. and 2.4. Results suggested that silica magnetic beads (no prior ultracentrifugation) and precipitation (without PCI/CI or ultrafiltration steps) recovered 670.12 ± 130.81 and 697.34 ± 144.55 µg DNA, respectively, accounting for roughly 33–35 % of input DNA (2 µg) and approximately 17–26 % when additional steps such as ultrafiltration and PCI/CI steps were included (see Supplementary Information, 3. for further details). Taking these isolation limitations into consideration, we note that the total DNA recovery from the minerals appears to likely be constrained not only by the lysis buffer but also to a considerable degree by the isolation approach.

### 3.2. Fragment range recovery from minerals

Resolving the DNA extracts on an agarose gel provided more compelling details of the observed differences described above and revealed that lysis buffers selectively retrieved bound DNA fragments from the minerals (Figure 2A-C). As expected, all silica magnetic beads extracts lacked fragments below 25 bp due to the cutoff introduced via the binding buffer while precipitation using LPA and ethanol recovered all fragments down to 10 bp.

**Figure 2 A-C.**
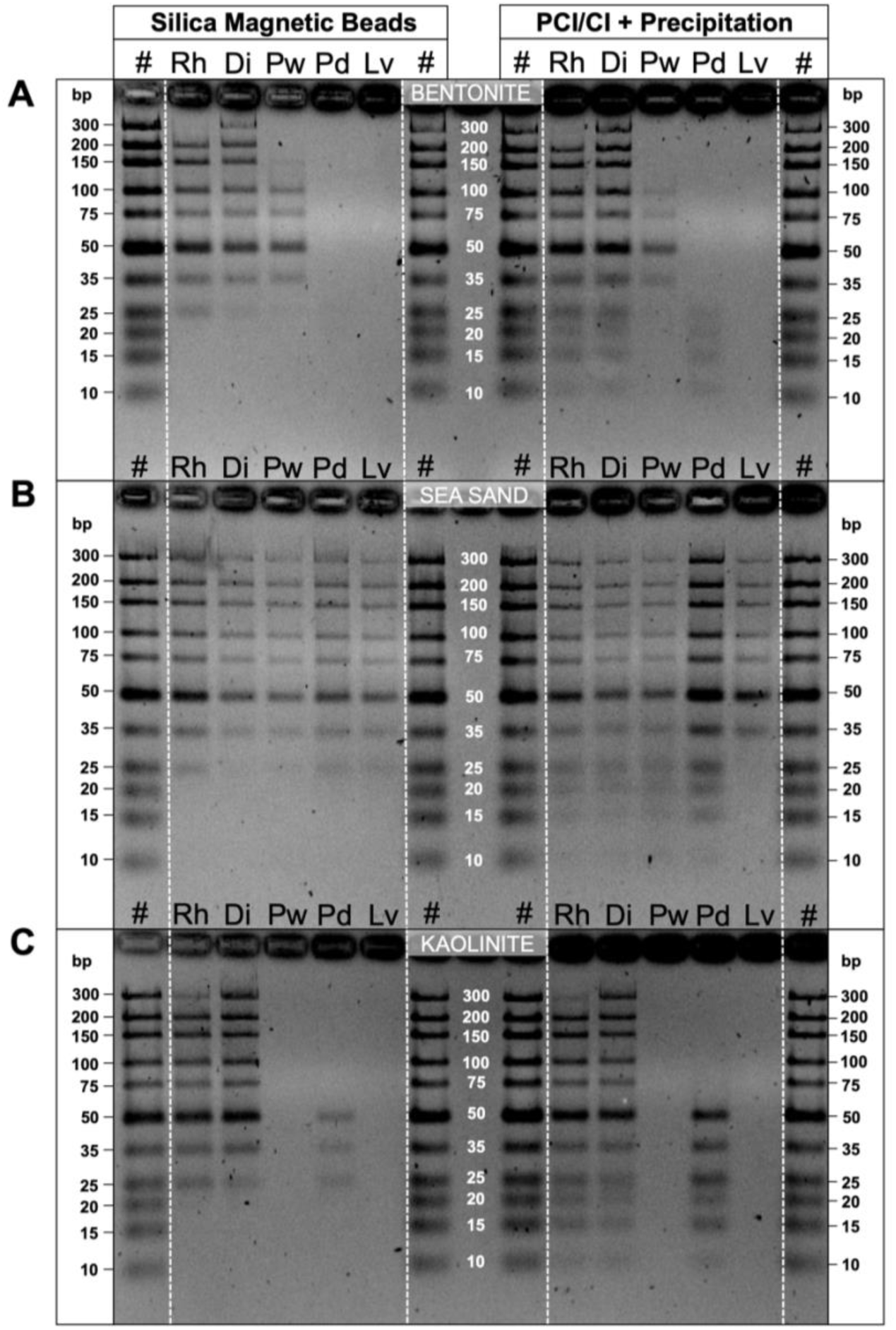
Gel-resolved mineral-desorbed DNA extracts. A) Bentonite, B) Sea-Sand, C) Kaolinite. The wells are labeled with the initials of the buffers used: (#) Ultra Low Range DNA marker, (Rh) Rohland buffer, (Di) Direito buffer, (Pw) PowerSoil buffer, (Pd) Pedersen buffer, (Lv) Lever buffer. Intensities of bands between the marker and the sample are comparable (∼ 500 ng). Silica magnetic beads fail to recover fragments below 25 bp in contrast to PCI/CI and precipitation isolation.

The differences in retrieved fragment ranges among the buffers broadly reflected the patterns observed in buffer performance on the basis of total recovered DNA. Rohland and Direito buffers recovered all fragments from bentonite with the exception of the very faint/missing 300 bp fragment when using the Rohland buffer (Figure 2A). Interestingly, the PowerSoil buffer was only able to desorb fragments in the 100 to 35 bp range, the Pedersen buffer only short fragments below and including 25 bp, and the Lever buffer no fragments at all. From kaolinite, Rohland and Direito buffers both recovered all fragments, Pedersen buffer again only short fragments – this time all below and including 50 bp, while with almost no DNA extracted, Lever and PowerSoil buffers did not show any visible bands (Figure 2C). In contrast to the two clay minerals, all buffers successfully recovered all fragments from sea-sand with the exception of the Lever buffer recovering fragments only down to 25 bp (Figure 2B).

These results further demonstrate that the buffer composition and lysis protocol itself not only directly impact the amount of desorbed DNA from the mineral, but also govern the recovery of different fragment lengths from mineral structures. While this is in line with previous findings by Slon et al. (2017), it expands on the initial observations discussed under 3.1. suggesting that lysis buffers containing high concentrations of EDTA (∼0.5M) or phosphate (∼1M) facilitate desorption of a wider range of DNA fragment sizes specifically from clay minerals compared to other buffers. Interestingly, lysis buffers with lower concentrations of EDTA such as Pedersen and Lever buffers performed better in recovery of DNA from sea-sand (quartz) compared to the clay minerals where they were able to desorb a narrower range of mostly shorter DNA fragments. This suggests that higher salt concentrations (i.e., high ionic strength) are required to desorb longer DNA fragments more effectively from clay minerals as these have been shown to be able to interact with more binding sites as opposed to shorter DNA fragments (Pietramellara et al. 2001).

In line with the highly fragmented nature of ancient DNA, this may suggest preferential use of “Bulat”-based buffers, such as the Pedersen buffer or PowerSoil buffer, for a more targeted desorption of shorter fragments and maximizing sequence yields in the lower fragment range most likely corresponding to ancient DNA. While this should be taken into consideration when extracting DNA from certain types of samples, extracting a broader suite of DNA fragment lengths might provide less bias in marine sediments where previous studies also reported successful recovery of 400-500 bp long 18S rDNA amplicons of microbial eukaryotes from Holocene deep-sea marine sediments (e.g., Boere et al. 2011a, Boere et al. 2011b, Boere et al. 2011c, Coolen et al. 2013).

Conversely, the choice of isolation approach could not have been shown to have a significant effect on total recovered DNA despite the length cut-off introduced by silica magnetic beads which would account for a 29.5% loss of the total DNA amount according to the contribution of fragments shorter than 25 bp in the ladder system.

Perhaps, the consequence of the length cut-off and the difference between the isolation approaches can be best illustrated in the case of DNA desorption from bentonite clay by the Pedersen buffer (Figure 2A). Here the length cut-off introduced by silica magnetic beads completely hinders the recovery of any DNA, leading to the premature conclusion that the lysate contains no DNA at all. In such cases, using the PCI/CI and precipitation approach over silica magnetic beads to retrieve all existing fragments could be of major importance in a first evaluation of the studied sample or area. Nevertheless, sequences below 30 bp are usually removed from bioinformatic analysis due to low mapping reliability (Shapiro and Hofreiter 2014, Allentoft et al. 2015), hence the cut-off below 25 bp introduced by silica magnetic beads can generally be considered advantageous for subsequent library preparation.

### 3.3. Sediment DNA extraction

Since Rohland and Direito buffers performed best, both in terms of total amount of recovered DNA and range of extracted DNA fragments from minerals (introducing the least bias among all buffers) we selected them for subsequent extraction of DNA from marine sediments.

Due to the depth/age of the chosen sediments, total recovered DNA from several sediment samples was below the limit of detection (Table SE1). Hence, sediment DNA extracts spiked with DNA ladder resolved on the gel allowed us to initially evaluate the performance of the two buffers and isolation approaches on natural environmental samples.

As expected, the performance of the two lysis buffers across the two isolation approaches was comparable to the results of the individual mineral types tested above. Both buffers retrieved DNA fragments for each of the sediment types in a comparable range (Figure S3). While short fragments (25 – 15 bp) did not bind efficiently to BS-SAP as suggested by their presence in the supernatant extracts after adsorption (data not shown), missing fragments in BS-LAC and SG-CLY extracts could point towards a stronger binding of short DNA fragments to components of the sediment matrix (Figure S3).

### 3.4. PCR inhibition of sediment extracts

Inhibition induced by co-extracted substances often presents a bottleneck in further downstream processing of sedimentary DNA extracts. Since we extracted DNA from deep marine sediments, we anticipated some degree of inhibition due to the presence of recalcitrant organic matter and the fact that the extracts exhibited strong coloration. Based on the results of the inhibition test (Figure S4), the Rohland buffer combined with silica magnetic beads produced the lowest number of inhibited extracts with only the extract from BS-SAP inhibiting the amplification reaction. This was anticipated due to the generally high organic content of the sapropel units (e.g., Bouloubassi et al. 1999). In contrast, the Direito buffer combined with silica magnetic beads yielded only a single non-inhibited extract (MID-PAC) which directly underlines the effect of lysis buffer on the extract quality. Surprisingly, DNA extracts of both buffers obtained by means of the PCI/CI and precipitation approach all showed inhibition (Figure S4). Since spiked extraction blanks for both buffers and isolation approaches showed the presence of amplified products, phenol residue is perhaps less likely reason for the inhibition but rather humic substances and other impurities coextracted with the DNA. Application of the One-Step Inhibitor Removal kit (Zymo Research Europe, Germany) greatly improved the quality of the previously inhibited extracts with the exception of BS-SAP and BS-LAC extracts of both lysis buffers isolated using the PCI/CI and precipitation approach that remained inhibited (Figure S5). Considering the difference in DNA extract quality between the two isolation approaches, not only are the silica magnetic beads more reliable in terms of extract quality but also in ease of work and safety by eliminating the exposure to hazardous chemicals and additional equipment. In addition, isolation using silica magnetic beads can be used either manually or deployed on automatic liquid handling systems to streamline the extraction and isolation of the DNA completely.

### 3.5. Sequencing library quality

Ideally, Illumina sequencing libraries exhibit a low or undetectable adapter dimer content and a high content of fragments of interest (fragments carrying informative inserts), reflected by high molarity ratios between the two regions as a measure for determining overall library quality.

Since ancient DNA is degraded, initial shearing of DNA is generally omitted in standard ancient DNA library preparation. However, the size range of fragments in the final library can still be broad depending on the sample preservation (e.g., modern contamination). Nevertheless, a right-skewed fragment distribution peaking at short insert sizes is indicative of degraded, ancient DNA.

In our analysis we constrained the range for fragments of interest to 157-527 bp as all libraries exhibited main fragment size peaks in this region, but we note the presence of longer fragments up to ∼1300 bp predominantly in SG-CLY libraries (404, 409 and 416) and library 414 (BS-SAP). We compare molarity ratios of libraries derived from sediment samples to the corresponding extraction blank of the respective combination of lysis buffer and isolation approach in order to infer overall quality and success of the preparation.

Initial comparisons suggest that libraries derived from the majority of sediment extracts exhibited notably higher molarity ratios than extraction blanks, indicating a higher concentration of aDNA inserts as opposed to library artifacts such as adapter dimers (Figure 3A). However, molarity ratios of sediment libraries appear highly heterogenous suggesting that a single combination of lysis buffer and isolation approach did not work optimally for all sediment types. On the one hand, Direito buffer combined with silica magnetic beads yielded libraries with the highest molarity ratios for N-PAC, MID-PAC, BS-SAP and SG-CLY sediments, while on the other hand, when combined with the PCI/CI and precipitation method the lowest molarity ratios for all sediment types were observed. This is also evident by the highest amount of amplification cycles that were required in the iPCR for this combination of lysis buffer and isolation approach compared to others (Table SE1). Moreover, BS-SAP, BS-LAC and SG-CLY molarity ratios were comparable to the corresponding blanks, suggesting unsuccessful library preparation likely due to remaining inhibition despite efforts to minimize it though extract dilution prior to library preparation. Unlike Direito, the Rohland lysis buffer exhibited higher molarity ratios and thus performed better when the PCI/CI and precipitation approach was used, but produced lower molarity ratios for certain extracts when combined with silica magnetic beads (Figure 3A).

**Figure 3 A-C.**
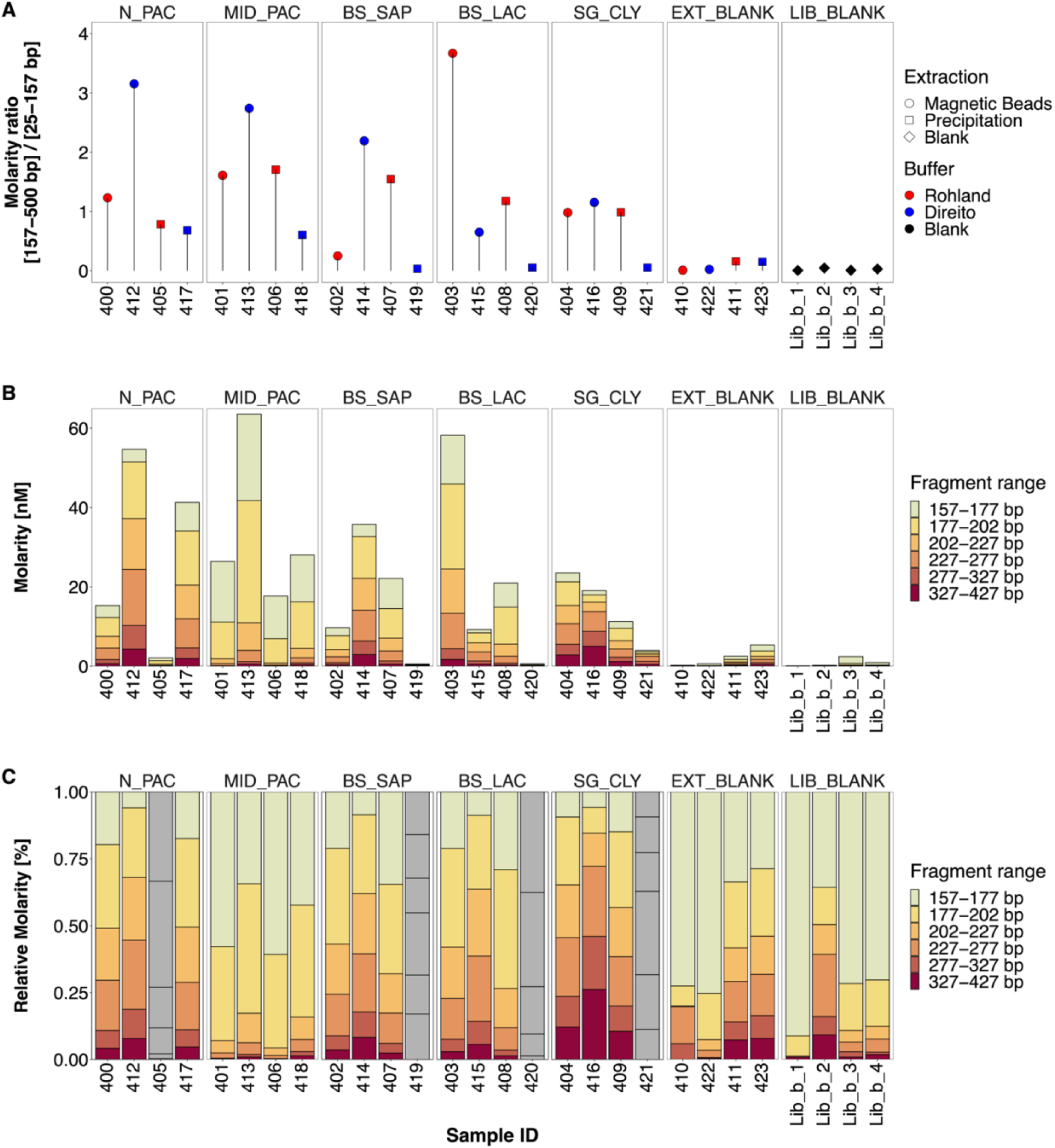
Characteristics of sequencing libraries prepared from sedimentary DNA extracts as inferred from a Fragment Analyzer. A) Molarity ratio between fragment ranges 157-500 bp and 25-157 bp, corresponding to aDNA inserts and adapter dimers, respectively. B) Absolute molarity of selected fragment ranges (nM). C) Relative molarity of selected fragment ranges (%) with grey bars corresponding to failed libraries. Samples are labelled by their Sample ID number (see Table S4).

Furthermore, absolute molarity of the combined fragment range (157-427 bp) closely followed the trends observed with the molarity ratios, suggesting that the amount of “sequencable” fragments largely defined the molarity ratio, with high adapter content being problematic only in certain sediment libraries. As already indicated above, several libraries exhibited a total molarity lower than those of the extraction blanks. We therefore considered libraries 405 (Rohland – PCI/CI and precipitation) and 419, 420, and 421 (Direito – PCI/CI and precipitation) as failed and excluded them from the further analysis and final interpretation (Figure 3B). While the total molarity of the 157-427 bp range varied between sediments and the four combinations of lysis buffer and isolation approach with no clear pattern, overall silica magnetic beads appear to have facilitated libraries with the highest total molarity compared to the PCI/CI and precipitation method, which in many cases failed. Since all sediment library preparations started with approximately similar amounts of DNA (Table SE1), these results accentuate the impact of inhibition on final library preparation success and quality derived from both the choice of lysis buffer and isolation approach. It is likely that the main source of inhibition can be attributed to co-extracted humic substances which have been shown to impact final library yield (Sidstedt et al. 2019). However, since we lack comprehensive details on organic matter content across different sediments, we cannot rule out potential other effects occurring during library preparation which are difficult to predict and detect.

While there seems to be no combination of buffer and isolation approach uniformly applicable to a wide array of marine sediment samples, we conclude that overall silica magnetic beads showed most reliability in facilitating successful library preparation.

### 3.6. Fragment size distribution

In contrast to the highly variable total molarity between the different combinations of lysis buffers and isolation approaches, relative molarity of selected fragment ranges appeared mostly consistent within each sediment. As expected, notable differences were mainly observed between different sediment types (Figure 3C). Still, in an effort to obtain a more comprehensive overview on global behavior of lysis buffers and isolation approaches and their effect on fragment size distribution, we further illustrated the absolute and relative molarity data grouped by fragment range treating sediments as replicates (Figure 4).

**Figure 4 A-B.**
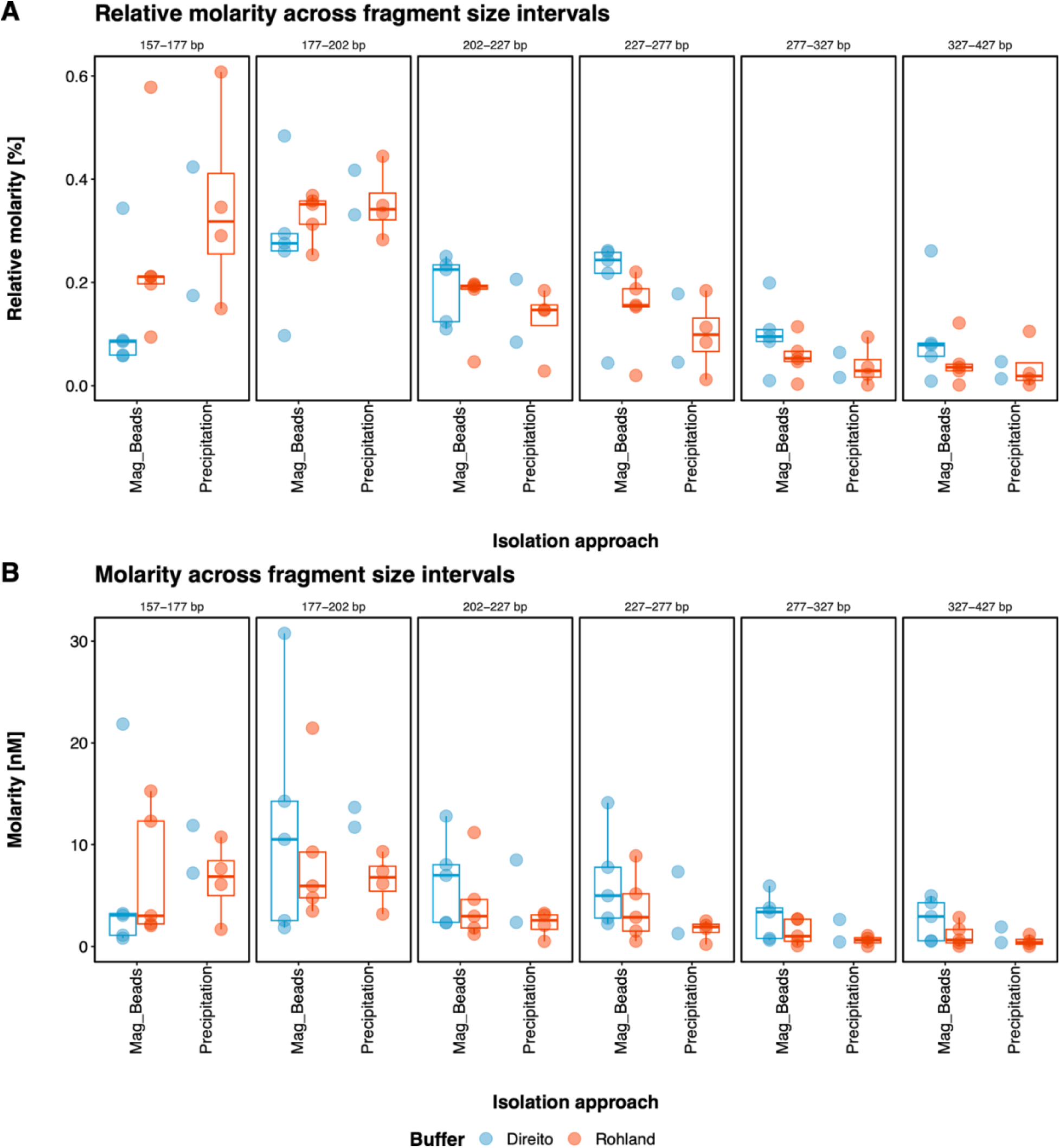
Relative (%) (A) and absolute molarity (nM) (B) of selected fragment ranges grouped by lysis buffer instead of sediment type (represented here as replicates). Libraries 405, 419, 420 and 421 were omitted from this analysis.

Here, the initial comparison of the two isolation methods indicates that silica magnetic beads exhibit slightly higher average absolute and relative molarity values in the longer fragment ranges (202-227 bp, 227-277 bp, 227-327 bp, 327-427 bp) and lower values in the shorter fragment ranges (157-177 bp and 177-202 bp) compared to the PCI/CI and precipitation approach (Figure 4). The inverse behavior of the two isolation methods could be connected to the fragment size cutoff introduced at 25 bp by the silica magnetic beads. However, since the lowest fragment size of 157 bp in this analysis would represent a ∼30 bp long insert, it is improbable that such a cutoff introduced by the beads during the extraction process explains this observation. Nevertheless, reaching final conclusions on definitive differences between the two isolation methods remains challenging and descriptive mainly due to inadequate replication in Direito-lysed samples due to 60% failed library preparations.

Strikingly however, we observed a notable difference between the lysis buffers across both isolation approaches where libraries derived from Direito buffer consistently appear to contain higher proportions of longer fragments and lower proportions of shorter fragments compared to the Rohland buffer derived libraries (Figure 4A). In an attempt to further corroborate these differences between lysis buffers, we ran independent linear mixed effects models for each fragment size range on relative molarities, accounting for differences between sediment types by introducing them as random factor. The analysis was conducted under the assumption that differences between the two isolation approaches were negligible in order to address the challenge of the limited number of replicates as discussed above. Under these conditions, the statistical analysis confirmed the above observations revealing significant differences between the two lysis buffers in longer fragment size ranges (**227-277 bp:** t = 3.421, df = 10.1, *P* = 0.0064; **277-327 bp:** t = 3.605, df = 10.1, *P* = 0.0047; **327-427 bp:** t = 2.995, df = 10.1, *P* = 0.0133) where higher proportions were detected in Direito buffer derived libraries, but also in the short fragment size range (**157-177 bp:** t = -4.634, df = 10.1, *P* = 0.0009), where proportions were higher in Rohland buffer derived libraries (Figure S6,A). Considering that the difference in the shorter range could be primarily a consequence of the non-independence of proportional values and that the difference in average absolute molarity between the lysis buffers in this fragment range was, in fact, opposite, we also repeated the statistical analysis with absolute molarity values. In this case, a significant difference between the lysis buffers was found only for the **327-427 bp** range (t = 2.584, df = 10.4, *P* = 0.0264; Figure S6,B).

Overall, our analysis of the sequencing libraries confirmed the effect of lysis buffer on fragment length distribution and suggests that phosphate-based lysis buffers such as Direito retrieve significantly more longer fragments compared to the EDTA-based lysis buffers such as Rohland. Similar observations on the impact of phosphate buffers on fragment size distribution has been previously reported by Ardelean et al. (2020) where a phosphate-based lysis buffer yielded longer reads compared to the Pedersen buffer in extracts from cave sediments. While based on our DNA-mineral desorption tests (Figure 2A,C) the difference between the phosphate-based Direito buffer and Pedersen buffer on sediment aeDNA extraction and subsequent library construction would presumably be more pronounced than the one we observed between Direito and Rohland buffers, results of Ardelean et al. (2020) nonetheless substantiate our observations. Furthermore, our DNA-mineral desorption tests revealed that the Rohland buffer exhibited a lower recovery of 300 bp long fragments from clay minerals in contrast to the Direito buffer, which may corroborate the observed lower retrieval in the 327-427 bp fragment range.

As mentioned by Ardelean et al. (2020), longer DNA fragments may originate from unknown active or dormant microorganisms. Considering the high abundance of microbial cells in marine subseafloor sediments estimated globally at 2.9·10^29^ cells (Kallmeyer et al. 2012) this is a likely possibility, but as mentioned before could also correspond to preserved and amplifiable fragments of eukaryotic DNA (e.g., Coolen et al. 2013). Although the taxonomic investigation of fragment sizes and final elucidation of the origin of longer fragments fall outside the scope of this study, we further emphasize on the importance of additional replication of the results presented above perhaps with a more extensive and diverse sample set to strengthen these conclusions.

## 4. Conclusions and recommendations

As our understanding of DNA preservation by means of mineral binding and its implications on reconstructing past ecosystems across temporal scales remains in continuous development, the results of this study illuminate capabilities of currently used sedimentary DNA extraction approaches for retrieving short DNA fragments bound to minerals and marine sediments.

Drawing from present comparisons and observations we demonstrated the profound impact of currently used lysis buffers and associated conditions on the amount of recovered mineral-bound DNA and corresponding fragment lengths. Depending on the study goals, sediment age, its location and mineralogical composition, the choice of lysis buffer should therefore be carefully considered.

Although we only tested five lysis conditions, we recommend the use of lysis buffers based on ∼0.5 M EDTA such as the Rohland buffer for aeDNA extractions from marine sediments as it proved the most reliable and versatile over various types of material (carbonaceous, siliceous, organic rich sapropel) and generally favored the recovery of short DNA fragments while still retrieving a wide range of fragment sizes.

However, further investigation and potential adaptation of other lysis buffers may provide improvements in their performance as well, specifically in regard to the complexity of marine sediments where DNA can interact with other naturally occurring minerals such as illite, chlorite, metal oxides or even complex organic matter that we have not tested in this study.

Consequently, initial screening and performance assessment of multiple buffers on target material could prove beneficial, specifically if the studied sediment consists of considerably different lithofacies where one buffer may provide a better compromise between DNA yield, fragment size distribution and inhibition than another.

Inhibition in general, and early inhibitor removal during DNA extraction in particular, should be considered vital to facilitate successful and optimal downstream processing of sedimentary DNA extracts while balancing cost effectiveness, safety and time investment. In this regard, silica magnetic beads took precedence over the classic liquid-phase organic extraction using phenol especially when combined with the Rohland lysis buffer, though at the small expense of a loss of fragments below 25 bp.

In conclusion, careful selection of DNA extraction approaches and their further optimization provides an important stepping stone for reliable recovery of mineral-bound DNA which in turn carry important implications for the ongoing development of aeDNA as a proxy for paleobiodiversity and paleoclimate in marine sciences and the field of sedimentary ancient DNA research in general.

## Acknowledgements

We thank Lasse Vinner for fruitful discussions and recommendations over lysis buffer and binding buffer composition adaptations. We thank the captains and the crew of RV *Meteor* Expedition M134, RV *Sonne* Expedition SO264 and RV *Poseidon* Expedition POS539 as well as David Aromokeye, Annika Schnakenberg and Jenny Wendt with the help in retrieving and cataloguing marine sediment samples used in this study. The work in this study was supported by the Research Centre Cluster of Excellence - EXC 2077 (project-ID 390741601) “The Ocean Floor – Earth’s Uncharted Interface” (MARUM), funded by the Deutsche Forschungsgemeinschaft (DFG) and the University of Bremen.

## Authors contributions

DG, MŽ and EW concieved the study. DG and CH designed the study. DG, CH & MŽ carried out experimental work. DG, CH and TRH carried out data analysis. MWF and EW supervised the study. DG wrote the manuscript with contributions from all authors.

## Conflict of interest statement

The authors declare no conflicts of interest.

## Data availability statement

The data that support the findings of this study and associated statistical analysis and plotting R scripts are openly available in Gande et al. (2023).

## Notes

https://zenodo.org/records/10318734

### Competing Interest Statement

The authors have declared no competing interest.

