## Supplementary information S1 for "Recovering short DNA fragments from minerals and marine sediments: a comparative study evaluating lysis and isolation approaches"

### Figures and Tables

**Figure S5 |** Gel resolved products of secondary 16S rRNA gene PCR inhibition test of previously inhibited sediment extracts that have been purified using the Zymo One-Step Inhibitor removal kit .. 10
**Figure S6 |** Relative (%) (A) and absolute molarity (nM) (B) recorded for selected fragment ranges in sequencing libraries of five tested sediment types ........ 11

#### 1. Modifications of the lysis conditions

To facilitate easier subsequent DNA isolation and better comparability between the used lysis conditions (Table 1), we introduced minor modifications to the protocols and adjusted the volume of lysis buffers as described below to fit a conventional 1.5-2.0 mL microcentrifuge tube for each step, enabling us to utilize standard laboratory equipment such as benchtop centrifuges. Briefly, we used double the amount of lysis buffer and 10 mM hexametaphosphate in “Lysis protocol II” by Lever et al. (2015) – 1.0 mL of “Cell lysis solution I” and 200 µL of 10 mM hexametaphosphate respectively. The volume of the lysis buffer described by Pedersen et al. (2016) was scaled down by a factor of three to 1.0 mL and supplemented with 5.75 µL 20 mg/mL proteinase K (Thermo Fischer Scientific, USA). In the protocol published by Epp et al. (2019) based on a modified Qiagen DNeasy Power-Max kit, we used 800 µL of CD1 lysis solution from Qiagen’s MagAttract PowerSoil kit (2020), added 5 µL of 20 mg/mL proteinase K (Thermo Fischer Scientific, USA) and then incubated samples for two hours at 45°C. After centrifugation, the supernatant was transferred to a fresh 2.0 mL tube and 200 µL of CD2 inhibitor removal solution from the same kit was added following the kit’s manual to obtain the final lysate. In the extraction protocol by Direito et al. (2012) we used 60 µL of CD1 solution from the Qiagen MagAttract PowerSoil kit instead of C1 solution originally called for, together with 1.0 mL of the lysis buffer. Lastly, in the lysis protocol published by Rohland et al. (2018), we did not pre-mix proteinase K with the buffer but added 12.5 µL of 20 mg/mL of the enzyme (Thermo Fischer Scientific, USA) directly to the 1.0 mL of lysis buffer previously added to the sample.

#### 2. Modifications of the protocol for isolation of DNA using silica magnetic beads

Isolation and purification of DNA using silica magnetic beads has previously been used in ancient DNA studies with a comprehensive protocol for sediment material targeting short ancient DNA fragments further refined by Rohland et al. (2018). The protocol proposes the use of *binding buffer D* (Dabney et al. 2013) for the successful binding of short DNA fragments to magnetic beads and recovery of fragments down to 35 bp. While working on a scale of a 2.0 mL microcentrifuge tube, however, this protocol utilizes only a fraction of the initial lysate (150 µL of 1000 µL). On the one hand, this is advantageous as the pure DNA extracts can be generated multiple times, but on the other hand, it can become limiting when the total DNA amount extracted from a sample is low or close to undetectable. In such cases, using the total lysate volume for DNA isolation would be beneficial. While it is possible to process multiple lysate subsamples in parallel using automatic liquid handling systems, manual extraction of many sub-samples in clean laboratory conditions is not feasible time- and material-wise. Neither it is ideal to use proportionally more binding buffer to lysate volume and work in larger tubes as such adaptations are usually not economical or available to each laboratory.

To circumvent these limitations, use the whole lysate and still constrain the DNA isolation to a 2.0 mL microcentrifuge tube, we up-concentrated the lysate after the extraction on an Amicon® Ultra 0.5 mL, 10 kDa Centrifugal Filter (MERCK, Germany) following an approach similar to Yang et al. (1998). This additional step not only allowed us to concentrate the lysate but also to wash away salts and (some) inhibitors.

With the ability to reduce the volume of the lysate from ~1.0 mL to ~100 µL we also assessed different binding buffer to lysate volumetric ratios to test if the ratio affects the retrieved fragment lengths and explored the possibility of using commercial binding buffers. For these tests, we again used GeneRuler Ultra Low DNA Ladder (Thermo Fischer Scientific, USA). Initially, we compared *binding buffer D* (Dabney et al. 2013, Rohland et al. 2018) and a modified Qiagen PB buffer (*BL01*) [500 mL PB, 2.5 mL 5M NaCl, 15 mL 3M Na-Acetate pH=5.2, 1 mL 37% HCl for final pH adjustment to 4-5]. Albeit with minor differences in volumes of added NaCl and Na-Acetate (L. Vinner, personal communication), *BL01* binding buffer was originally developed and described by Allentoft et al. (2015) where the authors have reported its better performance in binding shorter DNA fragments to silica in solution and was later also applied in the double stranded library preparation protocol by Meyer and Kircher (2010) on Qiagen's MinElute columns (L. Vinner, personal communication; Pedersen et al. 2016). We noticed that the recovery of fragments as low as 35 bp already occurs at a binding buffer to lysate volumetric ratio of 5:1 when using *BL01 buffer* compared to the *binding buffer D* at the suggested binding buffer to lysate volumetric ratio of 10:1 (1570 µL : 150 µL) (Rohland et al. 2018).

We then further tested other binding buffer to lysate ratios for *BL01* binding buffer – 10:1, 15:1 and 20:1 and resolved the DNA extracts on a 5% agarose gel. As shown in Figure S1, we observed that higher ratios - 15:1 and 20:1 of binding buffer to lysate recovered fragments down to 25 bp (indicated by stronger bands), ratio 10:1 showed a faint 25 bp band and 5:1 ratio no band at 25 bp at all, suggesting that higher ratios enable retrieval of shorter fragments – as similarly reported for *binding buffer G* (Glocke and Meyer 2017, Rohland et al. 2018) and in accordance with earlier observations by Allentoft et al. (2015).

Due to the convenience of easier preparation of the *BL01* binding buffer in contrast to *binding buffer D* or *G* and subsequent use of the same buffer in double-stranded library preparation (e.g., Pedersen et al. 2016), we selected *BL01* binding buffer at a volumetric ratio of 15:1 of binding buffer to lysate for isolation of DNA with silica magnetic beads for all extractions in this study.

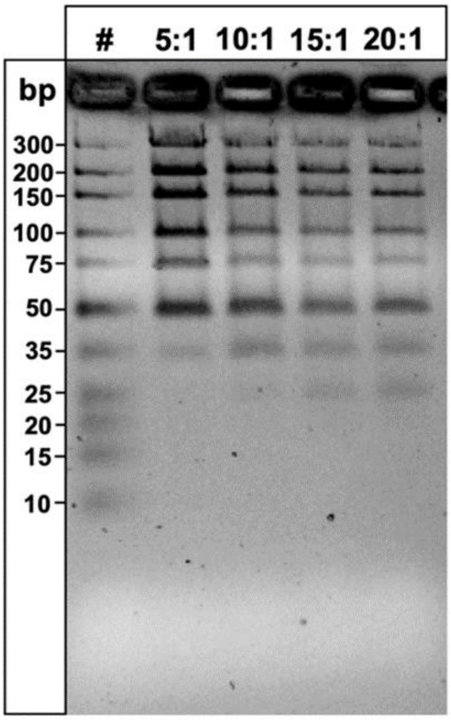

**Figure S1 |** Gel resolved fragments of Ultra Low DNA Ladder isolated using different volumetric ratios of *BL01* binding buffer to lysate. Marker well is labelled with (#) and the following wells with corresponding tested ratios.

##### 3. Unbound DNA and limits of the isolation method

We determined the amount of unbound DNA in the supernatants after the 24-hour adsorption period (Main text section 2.1) across the same two isolation approaches described in main text under 2.3 and 2.4. with an exception in the latter approach where we skipped the PCI/CI step and final cold ethanol wash. We used four replicates for each mineral and isolation approach. As shown in Table S1, the amount of unbound DNA was low on average with 0.39 % / 0.52 % (bentonite), 1.51 % / 0.77 % (sea-sand), 0.39 % / 0.26 % (kaolinite) either isolated with silica magnetic beads or the precipitation approach respectively.

As almost all DNA had bound to the mineral matrix, we further investigated to what degree DNA is lost during both isolation approaches or if it simply remains unextracted.

**Table S1 |** Total amount of unbound DNA (ng) recovered by the two isolation approaches

| n = 4 | Silica Magnetic Beads | PCI + CI / Precipitation |
|---|---|---|
| | **μ ± σ** | **μ ± σ** |
| **Bentonite** | 7.80 ± 5.20 | 10.38 ± 0.15 |
| **Sea-sand** | 30.24 ± 12.74 | 15.37 ± 2.75 |
| **Kaolinite** | 7.74 ± 5.16 | 5.23 ± 6.04 |

Protocol for isolation using silica magnetic beads (Main text, 2.3.) was tested in three stages using 4 µL (2 µg) of GeneRuler Ultra Low DNA Ladder (Thermo Fischer Scientific, USA) as following: *"Mag_Beads_only"* – DNA isolated from 120 µL lysate (4 µL DNA ladder + 116 µL EB buffer), *"Amicon + Mag_Beads"* – 1.0 mL lysate (4 µL DNA ladder + 996 µL EB buffer) first fully concentrated to 120 µL using Amicon® Ultra 0.5 mL, 10 kDa Centrifugal Filter (MERCK, Germany) and then isolated using beads and *"24h in ASW + Amicon + Mag_Beads"* – 1.0 mL lysate (4 µL DNA ladder + 996 µL artificial sea water) incubated in a 2.0 mL screw-cap tube for 24 hours (Main text, 2.1.), fully concentrated using Amicon® Ultra 0.5 mL, 10 kDa Centrifugal Filter (MERCK, Germany) to a volume of 120 µL and then isolated using beads.

We observed no loss of DNA due to possible binding to screw-cap tube walls and only a small reduction in total recovered DNA when the lysate was up-concentrated using Amicon® filters. However, silica magnetic beads accounted for the majority of non-recovered DNA with only 33.5 % of input DNA recovered on average (Figure S2).

Isolation with PCI/CI and precipitation (Main text, 2.4.) was also tested using 4 µL (2 µg) of GeneRuler Ultra Low DNA Ladder (Thermo Fischer Scientific, USA) in four stages as following: *"Precipitation"* – 500 µL lysate (4 µL DNA ladder + 496 µL EB buffer) mixed with LPA and ethanol, *"PCI + Precipitation"* – 1.0 mL lysate (4µL DNA ladder + 996 µL EB buffer) processed with PCI/CI, and DNA precipitated with LPA and ethanol, *"Amicon + Precipitation"* – 500 µL lysate (4 µL DNA ladder + 496 µL EB buffer) up-concentrated using Amicon® filters, volume brought back to 500 µL with EB buffer and DNA precipitated with LPA and ethanol and *"PCI + Amicon + Precipitation"* – 1.0 mL lysate (4µL DNA ladder + 996 µL EB buffer) processed with PCI/CI, up-concentrated using Amicon® filters, volume brought back to 500 µL with EB buffer and DNA precipitated with LPA and ethanol.

Recovery of DNA using precipitation was similar to isolation with silica magnetic beads with 34.88 % of input DNA recovered if precipitated directly. While no loss of DNA was observed when the PCI/CI step was included, Amicon® filters did account for additional loss of DNA compared to direct precipitation but recovered DNA in a similar range to when they were used with silica magnetic beads (Figure S2).

The low recovery of the two isolation approaches is surprising, specifically since silica magnetic beads have a binding capacity of > 4 mg DNA per mL of beads at a concentration of 50 mg/mL suggesting that 10 µL of the bead solution added for each DNA isolation should bind a maximum of 40 µg DNA.

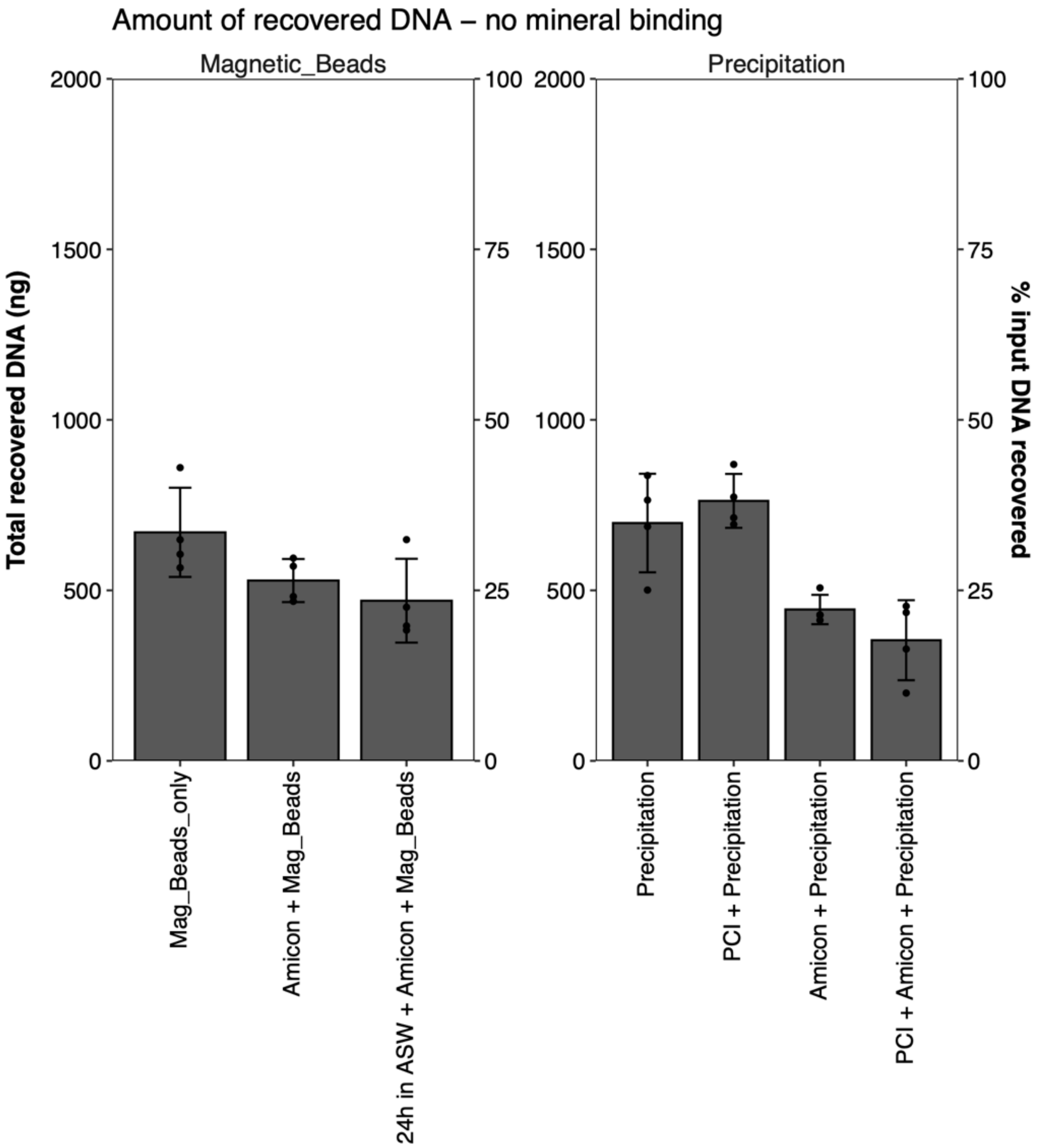

**Figure S2 |** Total recovered DNA (Ultra Low DNA ladder) through different stages of DNA isolation using silica magnetic beads or PCI + CI / precipitation. Input DNA was 2000 ng.

#### 4. Supporting figures and tables

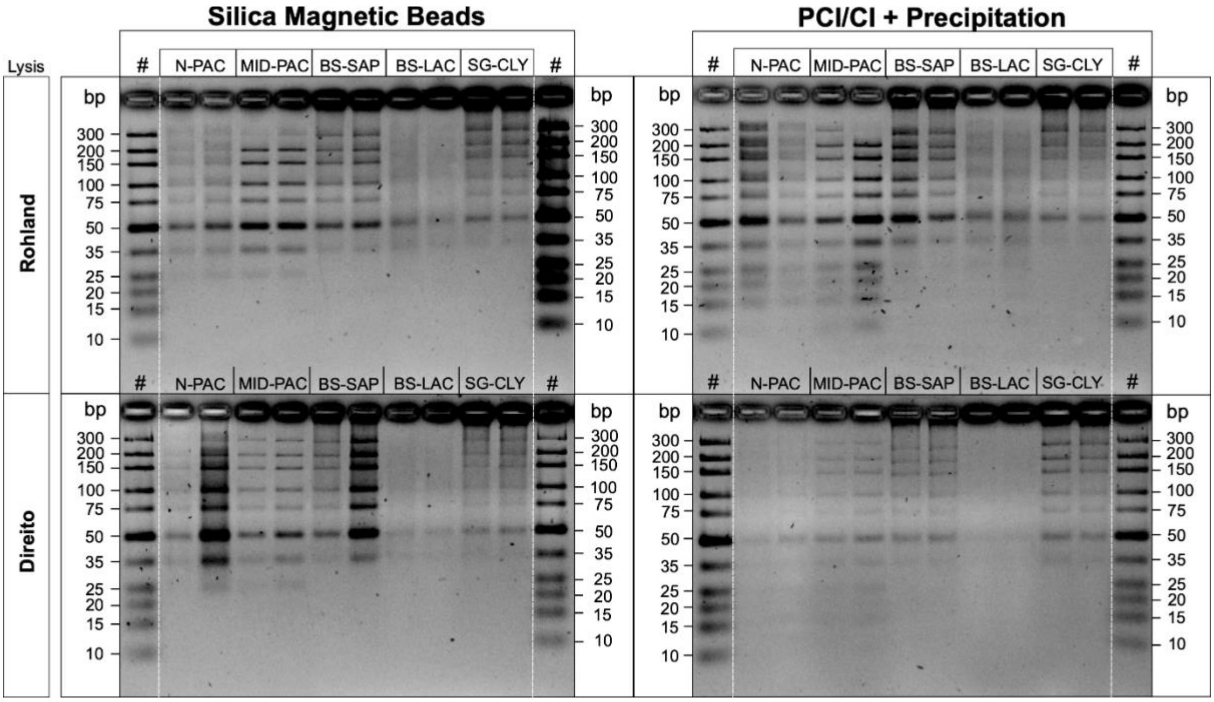

**Figure S3 |** Gel resolved DNA extracts from sediment samples spiked with 2 µg Ultra Low DNA ladder for the two lysis buffers and two isolation methods. Wells are labelled with sediment sample initials: (#) Ultra Low DNA ladder, (N-PAC) North-Pacific, (MID-PAC) Mid-Pacific, (BS-SAP) Black Sea sapropel unit, (BS-LAC) Black Sea lacustrine unit, (SG-CLY) South Georgia. Each sediment sample is represented in duplicates. Intensities of bands between the marker and the sample are comparable (~ 500 ng). Silica magnetic beads fail to recover fragments below 25 bp in contrast to precipitation.

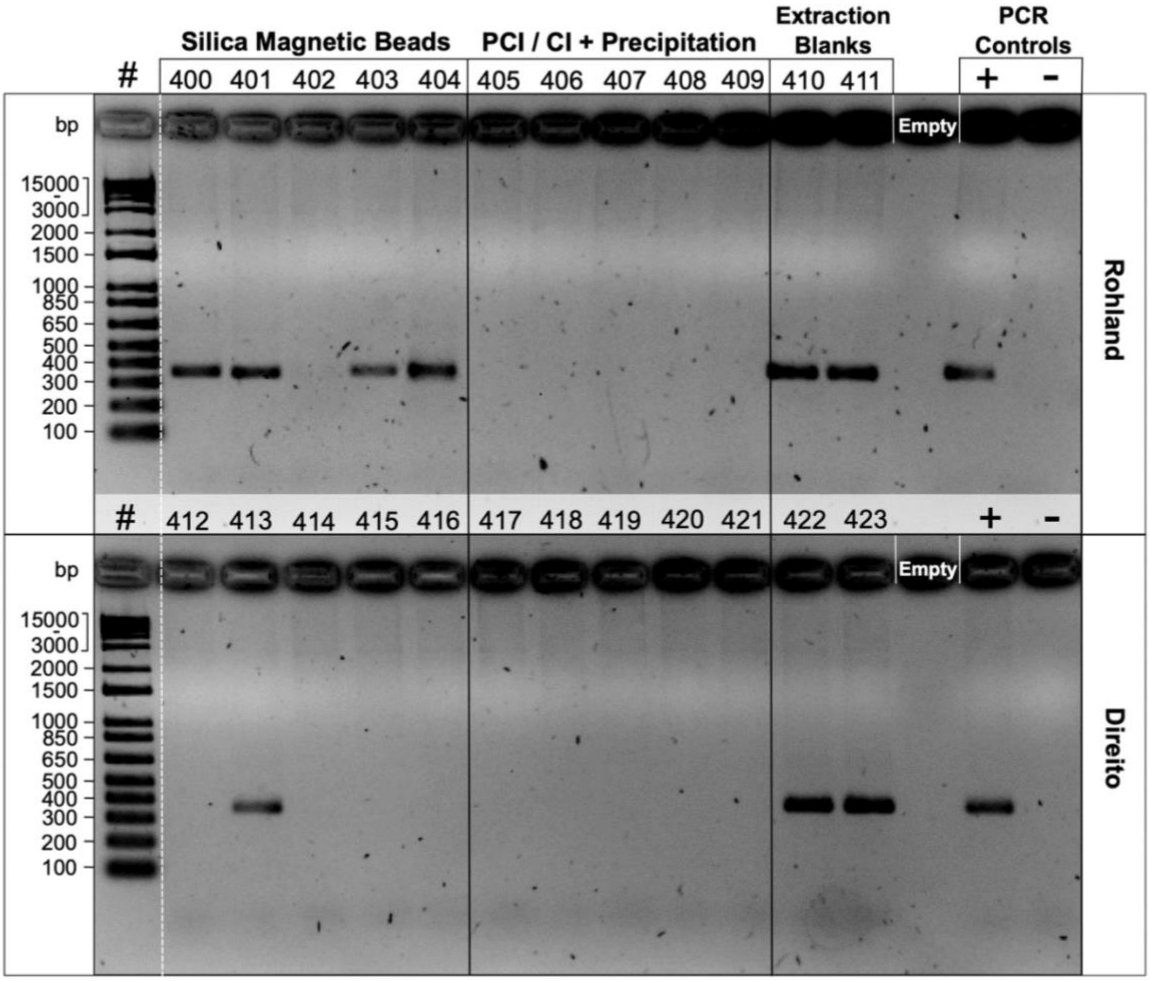

**Figure S4 |** Gel resolved products of the initial 16S rRNA gene PCR inhibition test. Marker well is labelled with (#), positive PCR control (+) and negative PCR control (-). Sample wells are labelled with a sample ID referring to the pooled replicate of the selected sediment sample (see Table S4). Absence of the 330 bp product represents an inhibited sample.

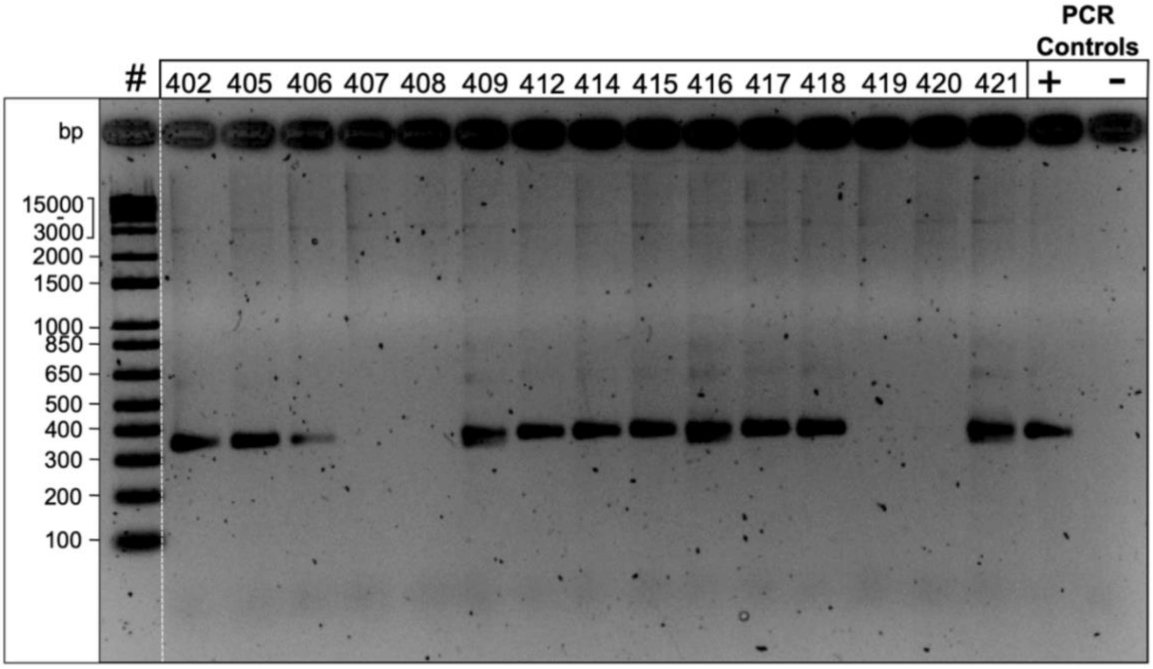

**Figure S5 |** Gel resolved products of secondary 16S rRNA gene PCR inhibition test of previously inhibited sediment extracts that have been purified using the Zymo One-Step Inhibitor removal kit. Marker well is labelled with (#), positive PCR control (+) and negative PCR control (-). Sample wells are labelled with a sample ID referring to the pooled replicate of the selected sediment sample (see Table S4). Absence of the 330 bp product represents an inhibited sample.

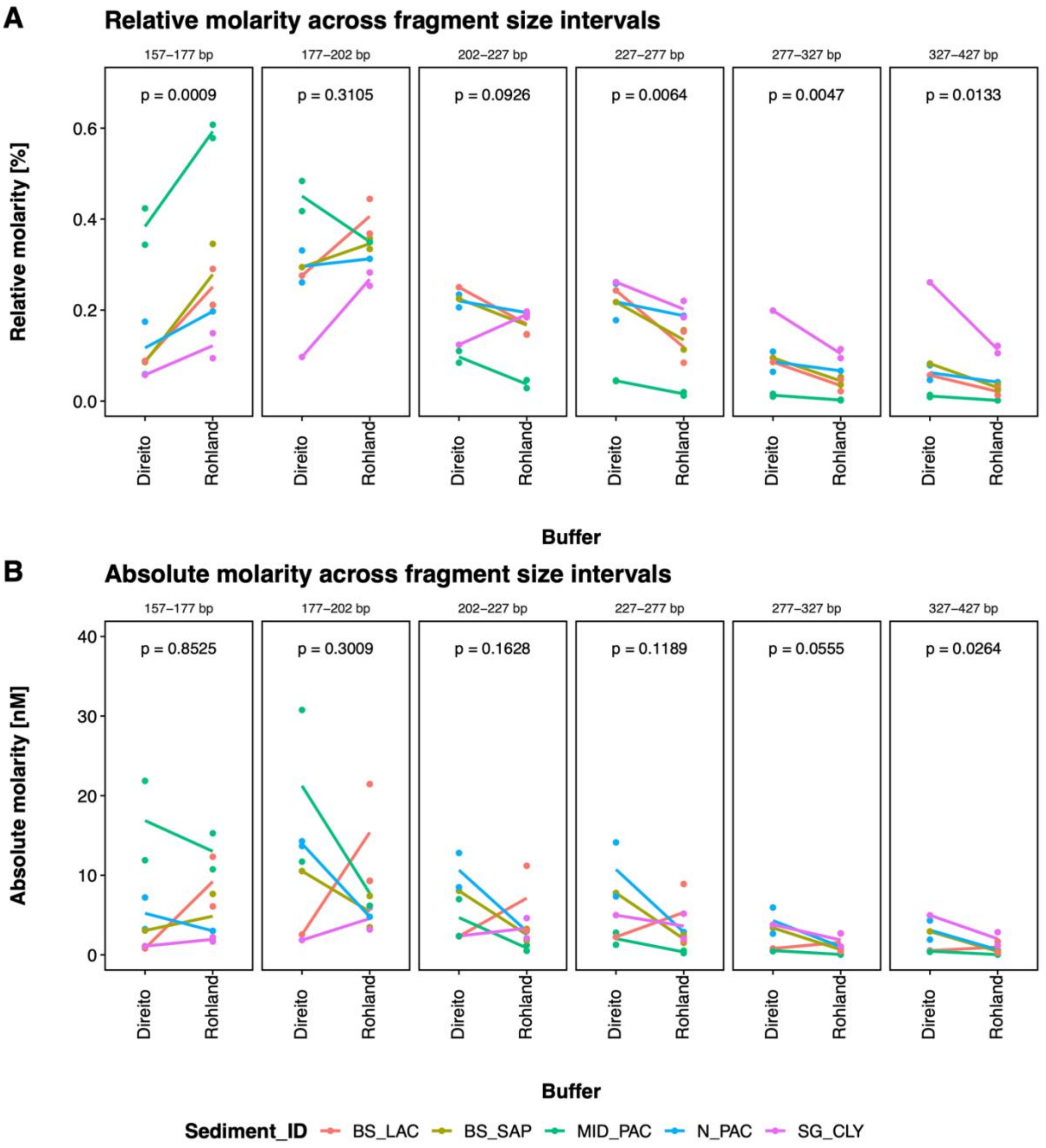

**Figure S6 |** Relative (%) (A) and absolute molarity (nM) (B) recorded for selected fragment ranges in sequencing libraries of five tested sediment types. The statistical significance of the difference between the libraries derived from sediment samples extracted with Direito and Rohland lysis buffers was assessed using general linear mixed effects models with sediment type as a random factor for each fragment range (p-values are provided in each panel).

**Table S2 |** Median (x̃), lower quartile (Q1), upper quartile (Q3) and interquartile range (IQR) values for total recovered DNA (ng) from the three minerals with five lysis buffers over two isolation approaches. Input DNA was 2000 ng.

| Isolation | Mineral | Buffer | x̃ | Q1 | Q3 | IQR |
|---|---|---|---|---|---|---|
| Silica Magnetic Beads | Bentonite | Rohland | 243.87 | 233.53 | 257.24 | 23.71 |
| | | Direito | 264.53 | 263.60 | 275.04 | 11.45 |
| | | PowerSoil | 120.87 | 67.58 | 197.94 | 130.36 |
| | | Pedersen | 15.44 | 14.89 | 15.52 | 0.63 |
| | | Lever | 0.00 | 0.0 | 1.57 | 1.57 |
| | Sea sand | Rohland | 249.88 | 243.68 | 255.51 | 11.84 |
| | | Direito | 192.37 | 175.69 | 202.58 | 26.88 |
| | | PowerSoil | 102.14 | 86.28 | 129.09 | 42.81 |
| | | Pedersen | 277.60 | 268.51 | 284.01 | 15.50 |
| | | Lever | 142.37 | 136.4 | 149.70 | 13.30 |
| | Kaolinite | Rohland | 328.40 | 321.88 | 339.28 | 17.40 |
| | | Direito | 352.14 | 306.39 | 397.89 | 91.50 |
| | | PowerSoil | 0.00 | 0.00 | 1.65 | 1.65 |
| | | Pedersen | 40.70 | 34.60 | 44.70 | 10.09 |
| | | Lever | 0.00 | 0.0 | 1.53 | 1.53 |
| PCI + CI / Precipitation | Bentonite | Rohland | 298.65 | 269.38 | 317.25 | 47.87 |
| | | Direito | 243.28 | 220.18 | 272.00 | 51.82 |
| | | PowerSoil | 53.08 | 49.7 | 74.10 | 24.39 |
| | | Pedersen | 39.32 | 32.68 | 48.57 | 15.89 |
| | | Lever | 7.78 | 7.67 | 8.17 | 0.49 |
| | Sea sand | Rohland | 253.55 | 225.38 | 274.16 | 48.78 |
| | | Direito | 131.54 | 110.75 | 156.65 | 45.90 |
| | | PowerSoil | 106.84 | 103.5 | 110.56 | 7.05 |
| | | Pedersen | 297.70 | 208.38 | 396.40 | 188.03 |
| | | Lever | 131.50 | 109.78 | 218.02 | 108.24 |
| | Kaolinite | Rohland | 293.41 | 273.06 | 323.43 | 50.37 |
| | | Direito | 292.93 | 232.60 | 313.09 | 80.49 |
| | | PowerSoil | 0.00 | 0.0 | 1.85 | 1.85 |
| | | Pedersen | 182.30 | 163.77 | 187.70 | 23.92 |
| | | Lever | 7.47 | 5.46 | 7.69 | 2.23 |

**Table S3 |** Results of non-parametric two-way Scheierer-Ray-Hare test on total recovered DNA from the three minerals

| | **Bentonite** | | | | **Sea-sand** | | | | **Kaolinite** | | | |
|---|---|---|---|---|---|---|---|---|---|---|---|---|
| | **df** | **SS** | **H** | **p-value** | **df** | **SS** | **H** | **p-value** | **df** | **SS** | **H** | **p-value** |
| **Buffer** | 4 | 5536.4 | 33.563 | **< 0.01** | 4 | 5540.7 | 25.0711 | **< 0.01** | 4 | 8591.1 | 40.752 | **< 0.01** |
| **Isolation approach** | 1 | 9.7 | 0.059 | 0.80849 | 1 | 72.0 | 0.3260 | 0.56803 | 1 | 21.8 | 0.103 | 0.74789 |
| **Interaction (Buffer x Isolation approach)** | 4 | 297.1 | 1.801 | 0.77228 | 4 | 247.5 | 1.1198 | 0.89112 | 4 | 276.2 | 1.310 | 0.85964 |
| **Residuals** | 34 | 1255.0 | | | 41 | 5201.8 | | | 40 | 1440.9 | | |

**Table S4 |** Sample codes and sediment sample information

| Sample ID | Sediment ID | Sediment Location | Sample depth (cm bsf) | Sediment description | Lysis buffer | Isolation approach |
|---|---|---|---|---|---|---|
| **400** | N-PAC | 49°18,447'N 168°33,427'E | 1026 | Greenish clay – diatomaceous clay (siliciclastic siliceous sediments) | Rohland | Magnetic Beads |
| **401** | MID-PAC | 41°34,914'N 170°25,783'E | 1127 | Calcareous nanno ooze (calcareous sediments) | Rohland | Magnetic Beads |
| **402** | BS-SAP | 43°28,919'N 030°11,077'E | 52.5 | Organic sapropel | Rohland | Magnetic Beads |
| **403** | BS-LAC | 43°28,919'N 030°11,077'E | 514 | Organic poor terrigenous mud (limnic) | Rohland | Magnetic Beads |
| **404** | SG-CLY | 54°9,458'S 37°58,581'W | 800 | Homogenous dry clay (clayish, diatom rich sediments) | Rohland | Magnetic Beads |
| **405** | N-PAC | 49°18,447'N 168°33,427'E | 1026 | Greenish clay – diatomaceous clay (siliciclastic siliceous sediments) | Rohland | PCI/CI + Precipitation |
| **406** | MID-PAC | 41°34,914'N 170°25,783'E | 1127 | Calcareous nanno ooze (calcareous sediments) | Rohland | PCI/CI + Precipitation |
| **407** | BS-SAP | 43°28,919'N 030°11,077'E | 52.5 | Organic sapropel | Rohland | PCI/CI + Precipitation |
| **408** | BS-LAC | 43°28,919'N 030°11,077'E | 514 | Organic poor terrigenous mud (limnic) | Rohland | PCI/CI + Precipitation |
| **409** | SG-CLY | 54°9,458'S 37°58,581'W | 800 | Homogenous dry clay (clayish, diatom rich sediments) | Rohland | PCI/CI + Precipitation |
| **410** | BLANK | - | - | - | Rohland | Magnetic Beads |
| **411** | BLANK | - | - | - | Rohland | PCI/CI + Precipitation |
| **412** | N-PAC | 49°18,447'N 168°33,427'E | 1026 | Greenish clay – diatomaceous clay (siliciclastic siliceous sediments) | Direito | Magnetic Beads |
| **413** | MID-PAC | 41°34,914'N 170°25,783'E | 1127 | Calcareous nanno ooze (calcareous sediments) | Direito | Magnetic Beads |
| **414** | BS-SAP | 43°28,919'N 030°11,077'E | 52.5 | Organic sapropel | Direito | Magnetic Beads |
| **415** | BS-LAC | 43°28,919'N 030°11,077'E | 514 | Organic poor terrigenous mud (limnic) | Direito | Magnetic Beads |
| **416** | SG-CLY | 54°9,458'S 37°58,581'W | 800 | Homogenous dry clay (clayish, diatom rich sediments) | Direito | Magnetic Beads |
| **417** | N-PAC | 49°18,447'N 168°33,427'E | 1026 | Greenish clay – diatomaceous clay (siliciclastic siliceous sediments) | Direito | PCI/CI + Precipitation |
| **418** | MID-PAC | 41°34,914'N 170°25,783'E | 1127 | Calcareous nanno ooze (calcareous sediments) | Direito | PCI/CI + Precipitation |
| **419** | BS-SAP | 43°28,919'N 030°11,077'E | 52.5 | Organic sapropel | Direito | PCI/CI + Precipitation |
| **420** | BS-LAC | 43°28,919'N 030°11,077'E | 514 | Organic poor terrigenous mud (limnic) | Direito | PCI/CI + Precipitation |
| **421** | SG-CLY | 54°9,458'S 37°58,581'W | 800 | Homogenous dry clay (clayish, diatom rich sediments) | Direito | PCI/CI + Precipitation |
| **422** | BLANK | - | - | - | Direito | Magnetic Beads |
| **423** | BLANK | - | - | - | Direito | PCI/CI + Precipitation |
